# Ethanolic extract of *Ocimum sanctum* leaves reduced invasion and matrix metalloproteinase activity of head and neck cancer cell lines

**DOI:** 10.1101/591214

**Authors:** Kusumawadee Utispan, Nattisa Niyomtham, Boon-ek Yingyongnarongkul, Sittichai Koontongkaew

**Author notes:** Corresponding author, (KU).

## Abstract

Head and neck squamous cell carcinoma (HNSCC) has a yearly incidence of 600,000 cases worldwide with a low survival rate. *Ocimum sanctum* L. or *Ocimum tenuiflorum* L. (Holy basil; Tulsi in Hindi), is a traditional medicine herb that demonstrates numerous effects, including anti-oxidant, anti-microbial, and anti-tumor effects. The aim of this study was to evaluate the anti-invasive effect of *O. sanctum* leaf extract on HNSCC cell lines. Ethanolic extract of *O. sanctum* leaf (EEOS) was prepared and the phenolic compounds were identified using high-performance liquid chromatography-electrospray ionization-time of flight-mass spectrometry. Genetically-matched HNSCC cell lines derived from primary (HN30 and HN4) and metastatic sites (HN31 and HN12) from the same patient were used in this study. The EEOS cytotoxicity to the cell lines was determined using an MTT assay. The invasion and matrix metalloproteinase (MMP)-2 and -9 activity of EEOS-treated cells were tested using a modified Boyden chamber assay and zymography, respectively. We found that EEOS significantly inhibited the invasion and MMP-2 and MMP-9 activity of HN4 and HN12 cells, but not HN30 and HN31 cells. Rosmarinic acid, caffeic acid, and apigenin were detected in EEOS. Moreover, rosmarinic acid was found as the major phenolic compound. Therefore, EEOS exerted its anti-invasive effect on HNSCC cells by attenuating MMP activity.

## Introduction

Head and neck squamous cell carcinoma (HNSCC) originates in the epithelial cells of the mucosal linings of the oral cavity, oropharynx, larynx, or hypopharynx [1]. HNSCC has an incidence of 600,000 cases per year worldwide, with a 40–50% mortality rate [2]. Similar to other tumors, invasion and metastasis are the critical processes that indicate HNSCC aggressiveness [3]. Matrix metalloproteinase (MMPs) are the key enzymes involved in tumor invasion and metastasis. MMP-2 and MMP-9 destroy the basement membrane and degrade the extracellular matrix, promoting tumor invasion [4]. Although modern medicine has contributed to treating cancers by surgery, chemotherapy, and radiotherapy, these modalities have not significantly changed the survival rate over the past three decades [5]. Thus, more effective treatments for local and metastatic HNSCC are needed.

*Ocimum sanctum* Linn. or *Ocimum tenuiflorum* Linn., commonly known as Holy Basil in English or Tulsi in Indian language [6], is a highly potent medicinal herb that is native throughout the eastern tropical countries including Thailand [7, 8]. *O. sanctum* is primarily composed of phytochemicals [9]. The fresh leaves and stem contain several flavonoids and phenolic compounds. Phenolic compounds such as apigenin, rosmarinic acid, cirsilineol, cirsimaritin, isothymusin, and isothymonin have been detected in *O. sanctum* leaf extracts [10]. The phytochemicals in this plant varies depending on different growing, harvesting, extraction, and storage conditions [11]. The leaf extracts of *O. sanctum* have numerous medicinal effects such as anti-oxidant [12, 13], wound healing [14], anti-microbial [15] and anti-tumorigenic effects [11]. The ethanol extract of *O. sanctum* leaf (EEOS) has demonstrated anti-tumorigenic effects on several cancer types including gastric cancer [16], pancreatic cancer [17], non-small cell lung cancer [18], and lung cancers [19, 20]. EEOS exhibited a variety of therapeutic effects on tumor cells. EEOS decreased the expression of proteins involved in the proliferation, invasion, and angiogenesis of carcinogen-induced rat gastric carcinoma [16]. Moreover, EEOS inhibited cancer invasion and metastasis. EEOS reduced vascular endothelial growth factor production and MMP-9 activity in metastatic-induced NCI-H460 non-small cell lung cancer cells by inhibiting the phosphatidylinositol 3-kinase (PI3K)/Akt signaling pathway [18]. Similarly, when treated with EEOS, Lewis lung carcinoma cell MMP-9 activity was inhibited [20].

This evidence suggests that EEOS has a wide range of activity against tumor cells. However, there is no data concerning the effect of EEOS on HNSCC. We hypothesized that EEOS would reduce HNSCC cell invasion. Therefore, this study evaluated the toxicity and anti-invasive effectof EEOS on primary and metastatic HNSCC cell lines. Moreover, the chemical constituents of the EEOS were identified.

## Materials and methods

### Chemicals

Caffeic acid, rosmaric acid, and apigenin were purchased from Sigma-Aldrich (St. Louis, MO). Acetonitrile (HPLC grade), ethanol, methanol, and formic acid (all analytical grade) were purchased from Merck (Darmstadt, Germany).

### *O. sanctum* leaf collection

The *O. sanctum* leaves were collected in the Phra Pradaeng district, Samut Prakan province, in central Thailand, during the rainy season. The collected leaves were authenticated for taxonomic identification by The Forest Botany Division, Forest Herbarium-BKF, Thailand.

### *O. sanctum* leaf extraction

The *O. sanctum* leaves were dried at room temperature and pulverized using a grinder. The *O. sanctum* powder was soaked in 95% ethanol for two weeks at room temperature. The extract was filtered through a 0.45 nm filter paper and concentrated using a rotary vacuum evaporator (Rotovapor R-215, BUCHI Labortechnik AG, Switzerland). The viscous residue was dried in a vacuum oven at 40°C. The ethanolic extract of *O. sanctum* (EEOS) was stored as a powder at 4°C until used.

### Cell culture

Genetically-matched HNSCC cell lines derived from primary and metastatic sites from the same patient were provided by Professor Silvio Gutkind (Moores Cancer Center, Department of Pharmacology, UCSD, CA, USA). The HN30 and HN31 cells were obtained from primary pharynx lesions and lymph node metastases (T3N0M0), respectively. The HN4 and HN12 cells were obtained from primary tongue lesions and lymph node metastases (T4N1M0), respectively [21]. The cells were maintained in Dulbecco’s Modified Eagle’s Medium (DMEM) (Invitrogen, Carlsbad, CA) supplemented with 10% fetal bovine serum, 100 U/ml penicillin, 100 μg/ml streptomycin (Invitrogen), and an anti-fungal agent. The cells were cultured in a 37°C humidified 5% CO_2_ atmosphere. The cells were passaged with 0.25% trypsin-EDTA when 90–100% confluent. Only cultures with at least 95%cell viability were used in the experiments.

### MTT assay

Cells (2,000 cells/well/100μl) were seeded in 96-well plates and incubated in a 37°C humidified 5% CO_2_ atmosphere. The cells were treated with 0.05, 0.1, 0.2, 0.4,or 0.8 mg/ml EEOS diluted in growth medium. Cells in growth medium served as control. After a 72 h incubation, the amount of viable cells in each treatment group were determined using thiazolylblue tetrazolium bromide (MTT, Sigma). The medium was removed, 150 μl of fresh medium was added, followed by adding 50 μl/well of 2 mg/ml MTT solution. The plates were incubated for 4 h at 37°C in a 5% CO_2_ incubator. The precipitated formazan crystals were solubilized in DMSO (200 μl/well). The absorbance of the resulting solution was measured at 570 nm by a microplate reader (Tecan trading, Austria) and converted to percent viable cells compared with control. Cell viability (%) was determined as follows: cell viability (%) = (mean Abs570_treated cells_ – mean Abs570_blank_)/(mean Abs570_control cells_ – mean Abs570_blank_) × 100. Three independent experiments were performed.

### Invasion assay

To evaluate cell invasion, an in vitro assay for cell invasion through Matrigel was performed using a blind-well Boyden chemotaxis chamber (Neuro Probe, Gaithersburg, MD) as previously described [22]. Briefly, the upper surface of 13 μm pore polycarbonate filters (Fisher Scientific, Canada) was coated with Matrigel, a reconstituted basement membrane gel (Corning, Tewksbury, MA) and placed between the upper and lower well plates of a blind-well Boyden chemotaxis chamber. Growth medium was used as a source of chemoattractants in the lower chamber. HNSCC cells (8×10^4^ cells) were resuspended in 0.4 mg/ml EEOS diluted in DMEM containing 0.1% BSA and were seeded into the upper well of the chamber. Cells treated with DMEM containing 0.1% BSA served as control. After 5 h incubation in a 37°C and 5% CO_2_ atmosphere incubator, the non-migrating cells on the upper surface of the filter were wiped off with a cotton bud. The filters were fixed with 0.5% crystal violet in 25% methanol for 10 min. The invaded cells on the lower surface of the filters were counted under a microscope at 400× magnification. Cell counting was performed by two investigators. Five randomly selected fields were counted per filter in each group, and the counts were averaged. Three independent experiments were performed.

### Conditioned medium preparation and zymography

HNSCC cells (2×10^6^ cells) were cultured in 6-well plates and incubated at 37°C for 24 h. After incubation, the wells were washed with PBS and treated with 0.4 mg/ml EEOS diluted in DMEM containing 0.1% BSA for 48 h. Cells cultured in DMEM containing 0.1% BSA were used as control. Conditioned medium (CM) was collected and centrifuged at 1,000 g and 4°C for 10 min. The CM was stored at −80°C until used. Total protein in the CM was estimated using the Pierce™ BCA protein assay kit (Thermo Fisher Scientific, Waltham, MA).

MMP-2 and MMP-9 activity in the CM were measured using gelatin zymography as previously described [23]. Briefly, gelatin (bloom 300, Sigma) was added to a 10% acrylamide separating gel at a final concentration of 0.2%.Samples containing equal amounts of total protein were mixed with non-reducing sample buffer and added to the gel. Following electrophoresis, the gels were washed in 2.5% Triton X-100 for 30 min at 37°C. The gels were incubated at 37°C overnight in developing buffer. The gels were stained with 0.5%Coomassie blue G250 in a 30% methanol and 10% glacial acetic acid solution for 30 min and destained in the same solution without Coomassie blue. The gelatin-degrading enzymes were identified as clear bands against the blue background of the stained gel. Images of the stained gels were captured under illumination using a G:BOX gel documentation system (Syngene, Frederick, MD). The gelatinolytic bands were quantified using GeneTools software (Syngene, Frederick, MD). Three independent experiments were performed.

### HPLC-ESI-TOF-MS analysis of EEOS

The *O. sanctum* extract was chemically analyzed using high-performance liquid chromatography coupled with electrospray ionization-time of flight-mass spectrometry (HPLC-ESI-TOF-MS). EEOS (1 mg/ml) was filtered through a 0.45 μm membrane filter and injected into an HPLC system (UltiMate^®^ 3000 system, Thermo Fisher Scientific, Sunnyvale, CA). Caffeic acid, rosmarinic acid, and apigenin standards were prepared (10, 50, 100, 150, 200, and 250 μg/ml). HPLC was performed using a reverse phase column (Symmetry C18 analysis column, 2.1 mm × 150 mm, and 5 μm particle size). A gradient elution was performed with a mobile phase of 0.1% formic acid (Component A) and 0.1% acetonitrile (Component B). Elution was performed at a 0.3 ml/min flow rate. The injection volume was 5 μl and the column temperature was 40°C. The components that were separated by the HPLC system were subjected to mass to charge ratio (m/z) analysis using a ESI-TOF-Ms system. ESI-TOF-MS was performed using a time of flight mass spectrometer (micrOTOF-Q-II, Bruker Daltonics, Germany). The ESI system negative-ion mode was used to generate m/z in a range 50–1000. The optimized mass spectrometric conditions were gas temperature (200°C), drying gas flow rate (8 l/min), nebulizer gas pressure (2 bar), and the capillary potential was 3000 V. Quantitative determination of the EEOS phenolic components was performed using a standard calibration curve. The data was analyzed using DataAnalysis 4.0 software (Bruker Daltonics, Germany).

### Statistical analysis

The results are presented as means and standard error of the mean (SEM). Statistical analysis was performed using one-way ANOVA followed by the Tukey’s multiple comparisons test with Prism GraphPad 7.0 (GraphPad Software, La Jolla, CA). The significance level was set at 0.05.

## Results

### Cytotoxic assessment of EEOS on the HNSCC cell lines

The cytotoxic effect of EEOS on the HNSCC cell lines was evaluated using an MTT assay (Fig 1). EEOS (0.8 mg/ml) significantly decreased the HN30, HN31, HN4, and HN12 cell viability to approximately 40%, 53%, 52%, and 40%, respectively, of that of their controls (*P* < 0.05). Whereas, 0.05, 0.1, 0.2, and 0.4 mg/ml EEOS were non-toxic to the cell lines. Therefore, the HNSCC cell lines were treated with 0.4 mg/ml EEOS and their invasion and MMP-2 and-9 activity were evaluated.

**Fig 1.**
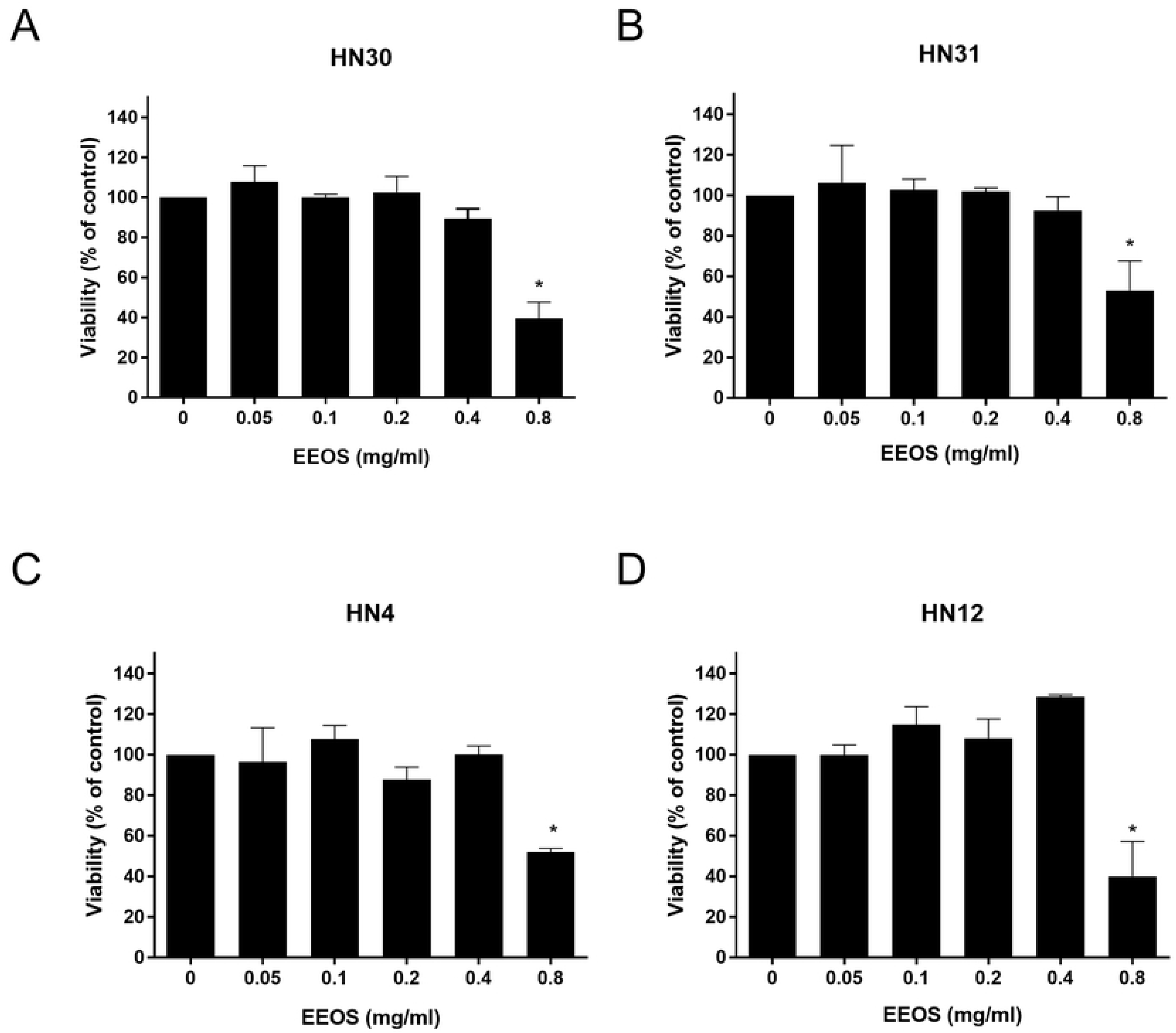
Cytotoxic evaluation of EEOS on HNSCC cells measured by MTT assay. EEOS in a range of concentrations were used to treat HN30 (A), HN31 (B), HN4 (C), and HN12 (D) cells for 72 h. Bars represent means±SEM (n=3). * indicates *p* < 0.05 compared with control.

### EEOS decreased metastatic HNSCC invasion

We found that HN30 and HN31 cell line invasion was not significantly inhibited by the non-toxic dose of EEOS (0.4 mg/ml) compared with control (*P* > 0.05) (Figs 2A and B). In contrast, 0.4 mg/ml EEOS significantly inhibited the invasion of the HN4 and HN12 cell lines by approximately 30% compared with control (*P* < 0.05) (Figs 2C and D).

**Fig 2.**
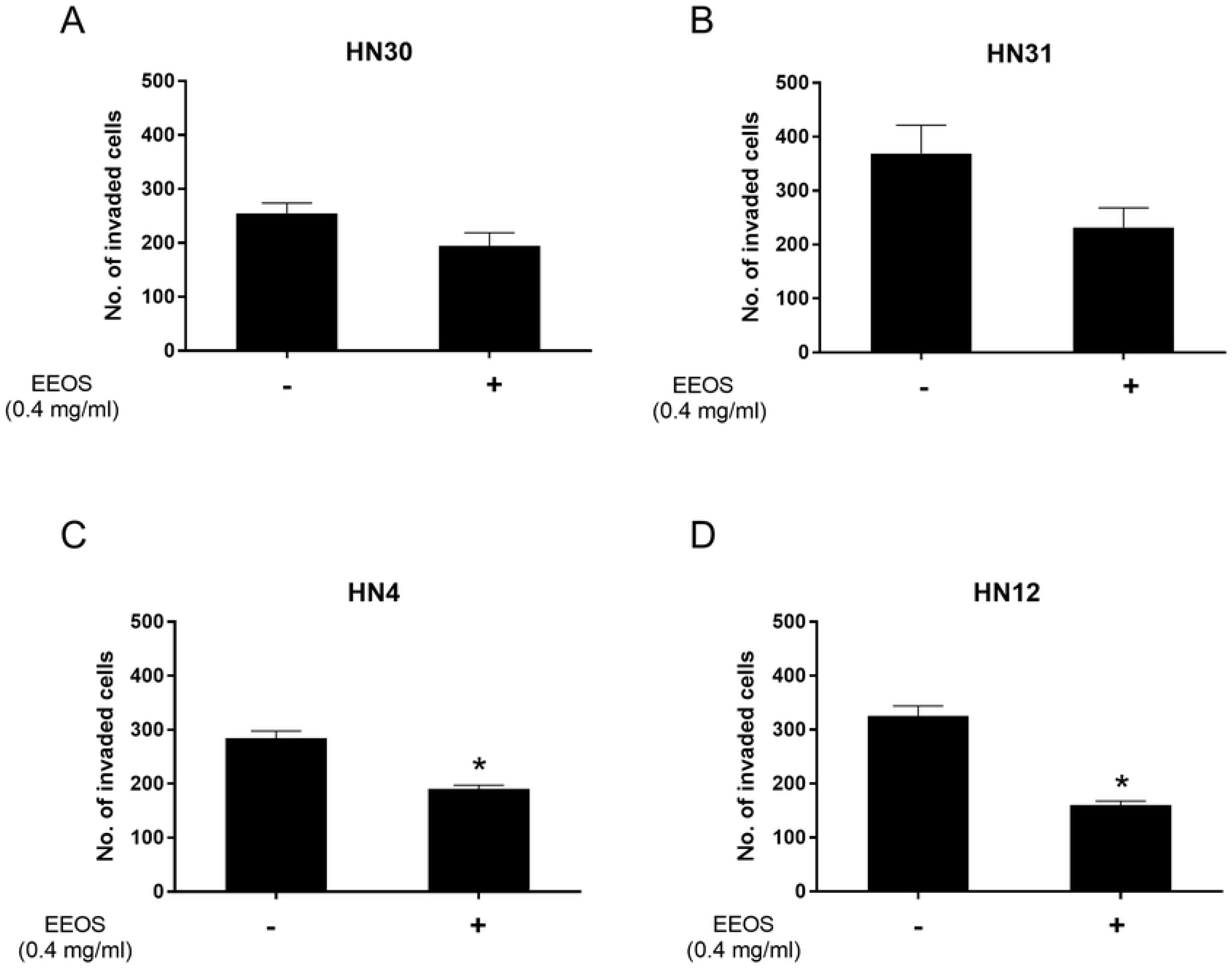
EEOS decreased HNSCC cell invasion. A non-cytotoxic dose of EEOS was used to treat HN30 (A), HN31 (B), HN4 (C), and HN12 (D) cells and evaluated cell invasion. Bars represent means±SEM (n=3). * indicates *p* < 0.05 compared with control.

### EEOS reduced MMP-2 and -9 activity of HNSCC cell lines

MMP-2 and MMP-9 activity was detected and quantified as gelatinolytic bands and arbitrary number of intensity, respectively. We found that 0.4 mg/ml EEOS treatment did not alter MMP-2 and MMP-9 activity of HN30 and HN31 cells (Fig 3A). However, MMP-2 and MMP-9 activity of HN4 and HN12 cells were downregulated when treated with EEOS (Fig 3B). Quantitative analysis of MMP activity revealed that the MMP-2 and MMP-9 activity in EEOS-treated HN30 and HN31 cells and control cells were not significantly different (*P* > 0.05) (Figs 3C and D). Differently, 0.4 mg/ml EEOS significantly reduced HN4 and HN12 cell MMP-2 activity to approximately 65% and 71%, respectively, of that of the control cells (*P* < 0.05). In addition, 0.4 mg/ml EEOS significantly reduced the MMP-9 activity of the HN4 and HN12 cells to approximately 44% and 85%, respectively, of that of the control cells (P < 0.05) (Figs 3C and D).

**Fig 3.**
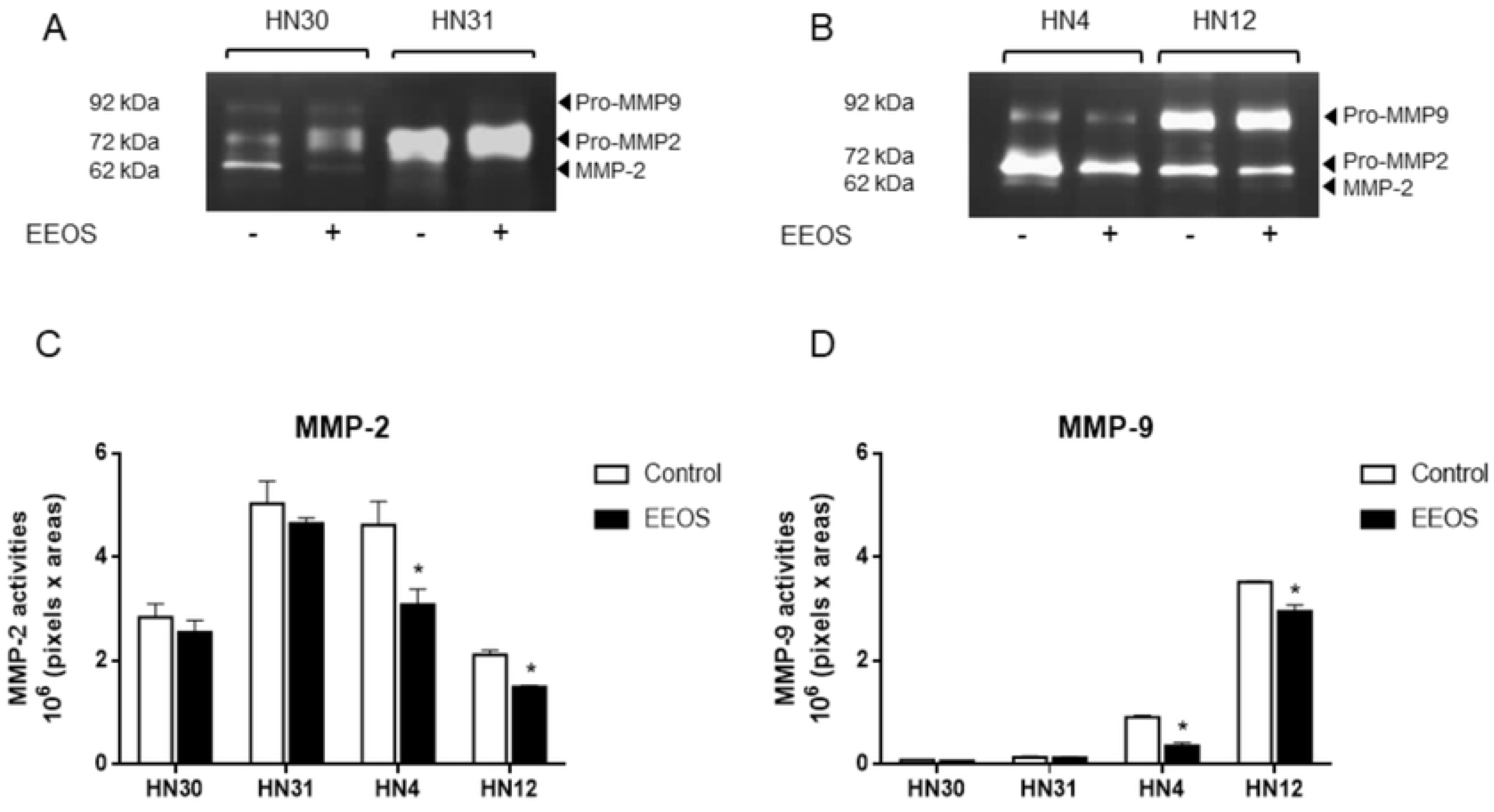
EEOS reduced MMP activity. The HNSCC cell lines were treated with 0.4 mg/ml EEOS for 48 h and the MMP activity in the conditioned media of HN30 and HN31 cells (A), and HN4 and HN12 cells (B) were detected using zymography. GeneTools software was used to quantify the gelatinolytic bands of MMP-2 (C) and MMP-9 (D) activity. Bars represent means±SEM (n=3). * indicates *p* < 0.05 compared with the control.

### HPLC analysis of EEOS

The HPLC retention times of caffeic acid, rosmarinic acid, and apigenin standards were determined (Fig 4A). The EEOS chromatograms demonstrated peaks 1, 2, and 3 with retention times that corresponded to those of the caffeic acid, rosmarinic acid, and apigenin standards, respectively (Fig 4B).

**Fig 4.**
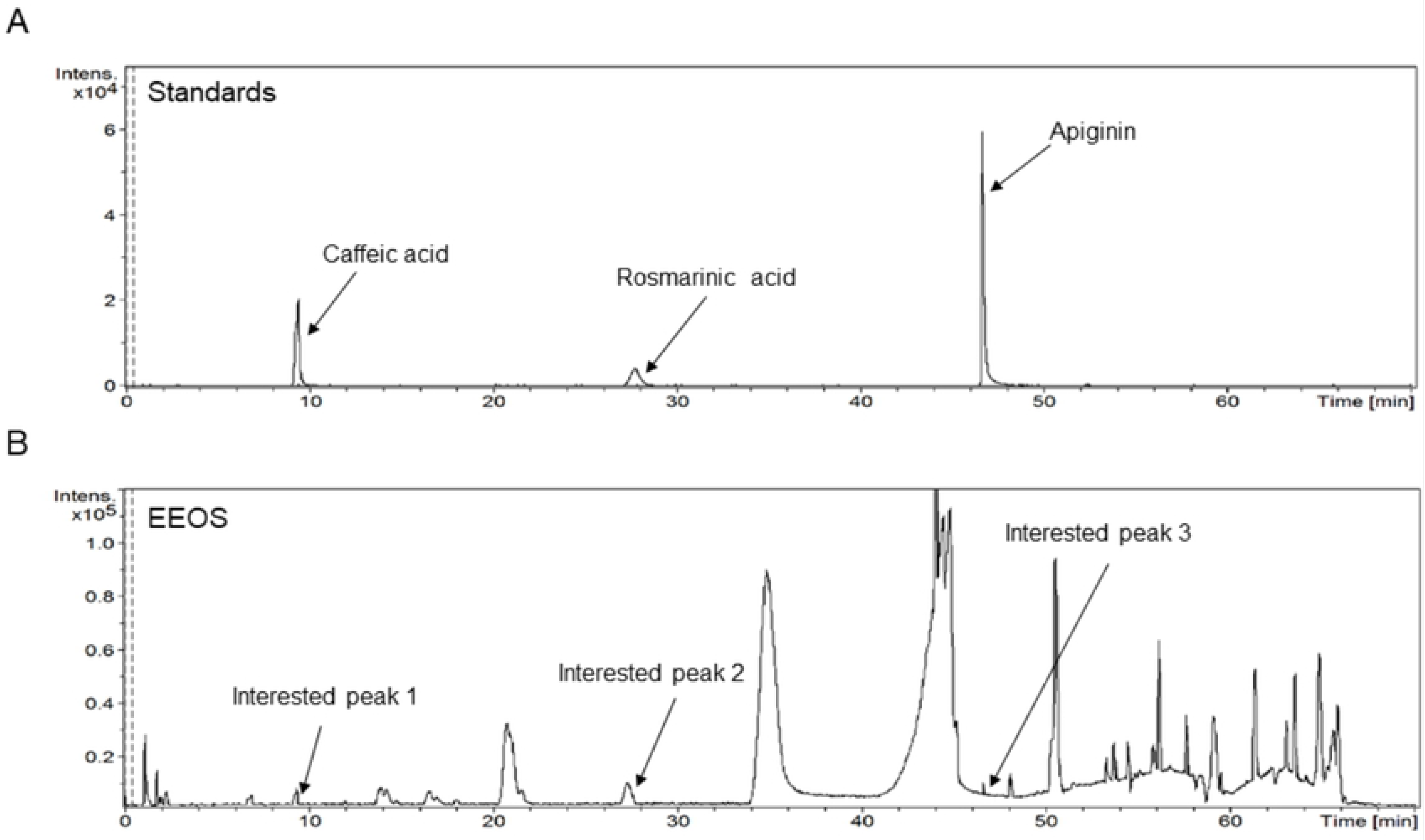
HPLC analysis of EEOS. Chromatogram of standard phenolic compounds (A) and the compounds detected in EEOS (B) are shown.

### Structural identification of the compounds in EEOS

HPLC-ESI-MS parameters were optimized and used to profile the EEOS. The selected 3 compounds in EEOS were putatively identified by comparison to the database (Table 1). Moreover, structures of the putative compounds were drawn by comparison to the known structure of standard compounds. The result revealed that compounds 1, 2, and 3 were caffeic acid, rosmarinic acid, and apigenin, respectively (Figs 5A and A). Quantification of the EEOS caffeic acid, rosmarinic acid, and apigenin indicated that rosmarinic acid was the major phenolic component (Fig 5C).

**Table 1.** HPLC-ESI-MS analysis of EEOS. Elucidation of empirical formulas and putative identification of each compound.

| Compounds | Putative identification | Retention time (min) | Empirical formula | Theoretical *m/z* | Experimental *m/z* |
| --- | --- | --- | --- | --- | --- |
| 1 | Caffeic acid | 9.40 | C<sub>9</sub>H<sub>8</sub>O<sub>4</sub> | 179.0349 | 179.0355 |
| 2 | Rosmarinic acid | 27.66 | C<sub>18</sub>H<sub>16</sub>O<sub>8</sub> | 359.0772 | 359.0764 |
| 3 | Apigenin | 46.63 | C<sub>15</sub>H<sub>10</sub>O<sub>5</sub> | 269.0455 | 269.0449 |

**Fig 5.**
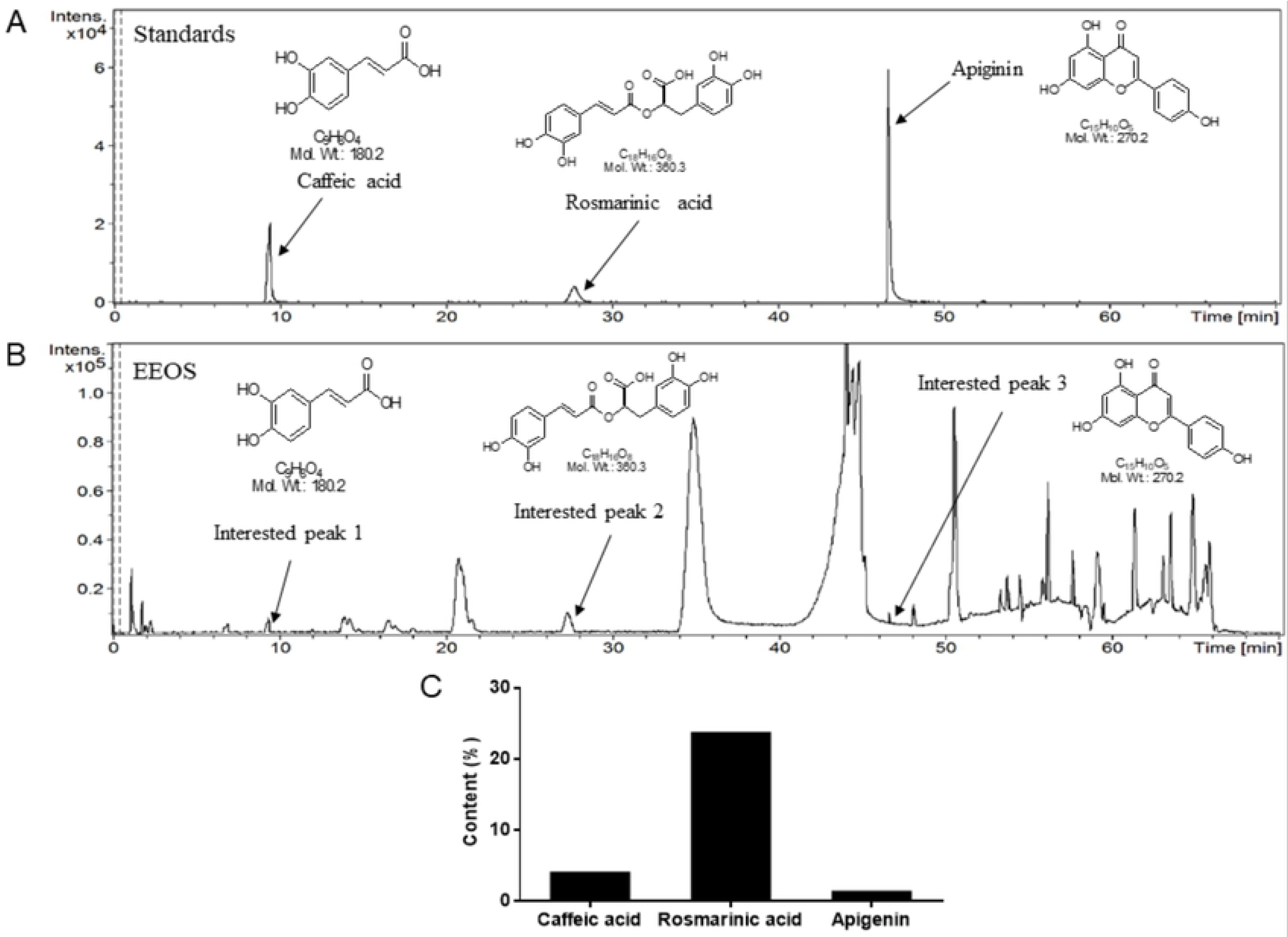
ESI-TOF-MS analysis of the compounds in EEOS. The structures of standard phenolic compounds (A) and the compounds in EEOS (B) analyzed from HPLC were identified. Quantification of the compounds in EEOS (C).

## Discussion

The present study investigated the effects of crude *O. sanctum* leaf extract on HNSCC invasiveness. EEOS cytotoxicity on the HNSCC cell lines was evaluated to determine the concentration to be used in subsequent experiments. A previous study reported that 0.2 mg/ml EEOS demonstrated a significant cytotoxic effect on a lung cancer cell line (A549) and mouse Lewis lung carcinoma cells [19]. However, 0.2 mg/ml EEOS was non-toxic to NCI-H460 non-small cell lung cancer cells [18]. Our findings indicated that 0.05, 0.1, 0.2, and 0.4 mg/ml EEOS did not reduce HNSCC cell viability whereas, 0.8 mg/ml EEOS was toxic to the cell lines evaluated. Thus, the cytotoxic effect of EEOS may be cell type-dependent.

MMP-2 and MMP-9 activity is important in initiating tumor invasion [4]. Several studies have reported the expression and role of MMP-2 and MMP-9 in HNSCC aggressiveness [24–27]. We evaluated the effect of EEOS on HNSCC invasion and MMP activity. We found that EEOS decreased HN4 and HN12 cell invasion by attenuating MMP-2 and MMP-9 activity. In contrast, there was no significant change in HN30 and HN31 cell invasion or MMP activity after EEOS treatment. Our results indicated that EEOS regulation of HNSCC invasiveness may be stage-dependent. These results imply that EEOS may have a potent role in inhibiting the invasiveness of a stage IV tumor. A study demonstrated that EEOS reduced the metastatic activity of Lewis lung carcinoma cell-injected mice [20]. They found that EEOS inhibited cancer invasion and MMP-9 activity, but not that of MMP-2. Another group confirmed that EEOS reduced MMP-9 and urokinase plasminogen activator activity in non-small cell lung cancer cells (NCI-H460) by inhibiting the PI3K/Akt signalling pathway [18]. MMP-9 production in HNSCC cell is induced through various signalling pathways, including epidermal growth factor receptor (EGFR), mitogen-activated kinase (MAPK) and PI3K/Akt [26, 28]. This suggests that EEOS may regulate MMP-2 and MMP-9 activity by targeting EGFR, MAPK, or PI3K/Akt pathways, leading to decreased HNSCC invasion.

The chemical composition of EEOS has been reported [29]. They found that EEOS was composed of several phenolic compounds and flavonoids, including rosmarinic acid and apigenin. Our results confirmed that the EEOS used in the present study contained rosmarinic acid, apigenin, and caffeic acid. Moreover, rosmarinic acid was the major phenolic component in our EEOS. Rosmarinic acid inhibited colon cancer invasion [30] colorectal cancer metastasis [31] by inhibiting MMP-2 and MMP-9 activity. These findings imply that the rosmarinic acid in EEOS may be a key factor in suppressing HNSCC cell invasion and MMP activity. Future studies should investigate the mechanism of rosmarinic acid and the other EEOS-derived components in suppressing HNSCC aggressiveness.

## Conclusions

Taken together, the present study demonstrated the cytotoxic and anti-tumorigenic effects of EEOS on HNSCC invasion and MMP-2 and MMP-9 activity. Interestingly, EEOS selectively regulated the invasion and MMP activity of HN4 and HN12 cells that were derived from stage IV tumors. We showed that the phenolic compounds rosmarinic acid, caffeic acid, and apigenin were present in EEOS. Moreover, rosmarinic acid was found as a major phenolic component. These results suggest that EEOS may be used as an alternative therapeutic agent in clinical research. However, the anti-tumorigenic mechanisms of the active compounds in EEOS require further investigation.

## Acknowledgements

The authors thank Professor Silvio Gutkind (Moores Cancer Center, Department of Pharmacology, UCSD, CA, USA) for the HNSCC cell lines used in our study. We thank Miss Hataichanok Yindeesompong, Miss Parncheewee Boonyawattananun, and Miss Nudda Khamrapich for technical assistance. We thank Dr. Amornmart Jaratrungtawee for HPLC-ESI-TOF-MS technical advice. The English editing of this manuscript was kindly performed by Dr. Kevin Tompkins, Office of Research Affairs, Faculty of Dentistry, Chulalongkorn University.

## References

1. Argiris A, Karamouzis MV, Raben D, Ferris RL. Head and neck cancer. Lancet. 2008;371(9625):1695–709. doi: 10.1016/S0140-6736(08)60728-X. PubMed PMID: 18486742.

2. Ferlay J, Soerjomataram I, Dikshit R, Eser S, Mathers C, Rebelo M, et al. Cancer incidence and mortality worldwide: sources, methods and major patterns in GLOBOCAN 2012. International journal of cancer. 2015;136(5):E359–86. doi: 10.1002/ijc.29210. PubMed PMID: 25220842.

3. Leemans CR, Snijders PJF, Brakenhoff RH. The molecular landscape of head and neck cancer. Nature reviews Cancer. 2018;18(5):269–82. doi: 10.1038/nrc.2018.11. PubMed PMID: 29497144.

4. Koontongkaew S. The tumor microenvironment contribution to development, growth, invasion and metastasis of head and neck squamous cell carcinomas. Journal of Cancer. 2013;4(1):66–83. doi: 10.7150/jca.5112. PubMed PMID: 23386906; PubMed Central PMCID: PMC3564248.

5. Kozakiewicz P, Grzybowska-Szatkowska L. Application of molecular targeted therapies in the treatment of head and neck squamous cell carcinoma. Oncology letters. 2018;15(5):7497–505. doi: 10.3892/ol.2018.8300. PubMed PMID: 29725456; PubMed Central PMCID: PMC5920345.

6. Baliga MS, Jimmy R, Thilakchand KR, Sunitha V, Bhat NR, Saldanha E, et al. Ocimum sanctum L (Holy Basil or Tulsi) and its phytochemicals in the prevention and treatment of cancer. Nutrition and cancer. 2013;65 Suppl 1:26–35. doi: 10.1080/01635581.2013.785010. PubMed PMID: 23682780.

7. Bast F, Rani P, Meena D. Chloroplast DNA phylogeography of holy basil (Ocimum tenuiflorum) in Indian subcontinent. TheScientificWorldJournal. 2014;2014:847482. doi: 10.1155/2014/847482. PubMed PMID: 24523650; PubMed Central PMCID: PMC3910118.

8. Suanarunsawat T, Anantasomboon G, Piewbang C. Anti-diabetic and anti-oxidative activity of fixed oil extracted from Ocimum sanctum L. leaves in diabetic rats. Experimental and therapeutic medicine. 2016;11(3):832–40. doi: 10.3892/etm.2016.2991. PubMed PMID: 26998000; PubMed Central PMCID: PMC4774317.

9. Pattanayak P, Behera P, Das D, Panda SK. Ocimum sanctum Linn. A reservoir plant for therapeutic applications: An overview. Pharmacognosy reviews. 2010;4(7):95–105. doi: 10.4103/0973-7847.65323. PubMed PMID: 22228948; PubMed Central PMCID: PMC3249909.

10. Gupta SK, Prakash J, Srivastava S. Validation of traditional claim of Tulsi, Ocimum sanctum Linn. as a medicinal plant. Indian journal of experimental biology. 2002;40(7):765–73. PubMed PMID: 12597545.

11. Bhattacharyya P, Bishayee A. Ocimum sanctum Linn. (Tulsi): an ethnomedicinal plant for the prevention and treatment of cancer. Anti-cancer drugs. 2013;24(7):659–66. doi: 10.1097/CAD.0b013e328361aca1. PubMed PMID: 23629478.

12. Kelm MA, Nair MG, Strasburg GM, DeWitt DL. Antioxidant and cyclooxygenase inhibitory phenolic compounds from Ocimum sanctum Linn. Phytomedicine : international journal of phytotherapy and phytopharmacology. 2000;7(1):7–13. doi: 10.1016/S0944-7113(00)80015-X. PubMed PMID: 10782484.

13. Manikandan P, Vidjaya Letchoumy P, Prathiba D, Nagini S. Combinatorial chemopreventive effect of Azadirachta indica and Ocimum sanctum on oxidant-antioxidant status, cell proliferation, apoptosis and angiogenesis in a rat forestomach carcinogenesis model. Singapore medical journal. 2008;49(10):814–22. PubMed PMID: 18946617.

14. Goel A, Kumar S, Singh DK, Bhatia AK. Wound healing potential of Ocimum sanctum Linn. with induction of tumor necrosis factor-alpha. Indian journal of experimental biology. 2010;48(4):402–6. PubMed PMID: 20726339.

15. Eswar P, Devaraj CG, Agarwal P. Anti-microbial Activity of Tulsi {Ocimum Sanctum (Linn.)} Extract on a Periodontal Pathogen in Human Dental Plaque: An Invitro Study. Journal of clinical and diagnostic research : JCDR. 2016;10(3):ZC53-6. doi: 10.7860/JCDR/2016/16214.7468. PubMed PMID: 27135002; PubMed Central PMCID: PMC4843387.

16. Manikandan P, Vidjaya Letchoumy P, Prathiba D, Nagini S. Proliferation, angiogenesis and apoptosis-associated proteins are molecular targets for chemoprevention of MNNG-induced gastric carcinogenesis by ethanolic Ocimum sanctum leaf extract. Singapore medical journal. 2007;48(7):645–51. PubMed PMID: 17609827.

17. Shimizu T, Torres MP, Chakraborty S, Souchek JJ, Rachagani S, Kaur S, et al. Holy Basil leaf extract decreases tumorigenicity and metastasis of aggressive human pancreatic cancer cells in vitro and in vivo: potential role in therapy. Cancer letters. 2013;336(2):270–80. doi: 10.1016/j.canlet.2013.03.017. PubMed PMID: 23523869; PubMed Central PMCID: PMC3700662.

18. Kwak TK, Sohn EJ, Kim S, Won G, Choi JU, Jeong K, et al. Inhibitory effect of ethanol extract of Ocimum sanctum on osteopontin mediated metastasis of NCI-H460 non-small cell lung cancer cells. BMC complementary and alternative medicine. 2014;14:419. doi: 10.1186/1472-6882-14-419. PubMed PMID: 25345853; PubMed Central PMCID: PMC4219006.

19. Magesh V, Lee JC, Ahn KS, Lee HJ, Lee HJ, Lee EO, et al. Ocimum sanctum induces apoptosis in A549 lung cancer cells and suppresses the in vivo growth of Lewis lung carcinoma cells. Phytotherapy research : PTR. 2009;23(10):1385–91. doi: 10.1002/ptr.2784. PubMed PMID: 19277950.

20. Kim SC, Magesh V, Jeong SJ, Lee HJ, Ahn KS, Lee HJ, et al. Ethanol extract of Ocimum sanctum exerts anti-metastatic activity through inactivation of matrix metalloproteinase-9 and enhancement of anti-oxidant enzymes. Food and chemical toxicology : an international journal published for the British Industrial Biological Research Association. 2010;48(6):1478–82. doi: 10.1016/j.fct.2010.03.014. PubMed PMID: 20233602.

21. Cardinali M, Pietraszkiewicz H, Ensley JF, Robbins KC. Tyrosine phosphorylation as a marker for aberrantly regulated growth-promoting pathways in cell lines derived from head and neck malignancies. International journal of cancer. 1995;61(1):98–103. PubMed PMID: 7705939.

22. Albini A, Iwamoto Y, Kleinman HK, Martin GR, Aaronson SA, Kozlowski JM, et al. A rapid in vitro assay for quantitating the invasive potential of tumor cells. Cancer research. 1987;47(12):3239–45. PubMed PMID: 2438036.

23. Thomas GJ, Lewis MP, Hart IR, Marshall JF, Speight PM. AlphaVbeta6 integrin promotes invasion of squamous carcinoma cells through up-regulation of matrix metalloproteinase-9. International journal of cancer. 2001;92(5):641–50. PubMed PMID: 11340566.

24. de Vicente JC, Fresno MF, Villalain L, Vega JA, Hernandez Vallejo G. Expression and clinical significance of matrix metalloproteinase-2 and matrix metalloproteinase-9 in oral squamous cell carcinoma. Oral oncology. 2005;41(3):283–93. doi: 10.1016/j.oraloncology.2004.08.013. PubMed PMID: 15743691.

25. Kato K, Hara A, Kuno T, Kitaori N, Huilan Z, Mori H, et al. Matrix metalloproteinases 2 and 9 in oral squamous cell carcinomas: manifestation and localization of their activity. Journal of cancer research and clinical oncology. 2005;131(6):340–6. doi: 10.1007/s00432-004-0654-8. PubMed PMID: 15614523.

26. Koontongkaew S, Amornphimoltham P, Monthanpisut P, Saensuk T, Leelakriangsak M. Fibroblasts and extracellular matrix differently modulate MMP activation by primary and metastatic head and neck cancer cells. Medical oncology. 2012;29(2):690–703. doi: 10.1007/s12032-011-9871-6. PubMed PMID: 21380786.

27. Patel BP, Shah SV, Shukla SN, Shah PM, Patel PS. Clinical significance of MMP-2 and MMP-9 in patients with oral cancer. Head & neck. 2007;29(6):564–72. doi: 10.1002/hed.20561. PubMed PMID: 17252594.

28. P Oc, Wongkajornsilp A, Rhys-Evans PH, Eccles SA. Signaling pathways required for matrix metalloproteinase-9 induction by betacellulin in head-and-neck squamous carcinoma cells. International journal of cancer. 2004;111(2):174–83. doi: 10.1002/ijc.20228. PubMed PMID: 15197768.

29. Venuprasad MP, Kandikattu HK, Razack S, Amruta N, Khanum F. Chemical composition of Ocimum sanctum by LC-ESI-MS/MS analysis and its protective effects against smoke induced lung and neuronal tissue damage in rats. Biomedicine & pharmacotherapy = Biomedecine & pharmacotherapie. 2017;91:1–12. doi: 10.1016/j.biopha.2017.04.011. PubMed PMID: 28433747.

30. Xu Y, Xu G, Liu L, Xu D, Liu J. Anti-invasion effect of rosmarinic acid via the extracellular signal-regulated kinase and oxidation-reduction pathway in Ls174-T cells. Journal of cellular biochemistry. 2010;111(2):370–9. doi: 10.1002/jcb.22708. PubMed PMID: 20506543.

31. Han YH, Kee JY, Hong SH. Rosmarinic Acid Activates AMPK to Inhibit Metastasis of Colorectal Cancer. Frontiers in pharmacology. 2018;9:68. doi: 10.3389/fphar.2018.00068. PubMed PMID: 29459827; PubMed Central PMCID: PMC5807338.

